# A likelihood-based framework for demographic inference from genealogical trees

**DOI:** 10.1101/2023.10.10.561787

**Authors:** Caoqi Fan, Jordan L. Cahoon, Bryan L. Dinh, Diego Ortega-Del Vecchyo, Christian Huber, Michael D. Edge, Nicholas Mancuso, Charleston W.K. Chiang

## Abstract

The demographic history of a population drives the pattern of genetic variation and is encoded in the gene-genealogical trees of the sampled alleles. However, existing methods to infer demographic history from genetic data tend to use relatively low-dimensional summaries of the genealogy, such as allele frequency spectra. As a step toward capturing more of the information encoded in the genome-wide sequence of genealogical trees, here we propose a novel framework called the genealogical likelihood (gLike), which derives the full likelihood of a genealogical tree under any hypothesized demographic history. Employing a graph-based structure, gLike summarizes across independent trees the relationships among all lineages in a tree with all possible trajectories of population memberships through time and efficiently computes the exact marginal probability under a parameterized demographic model. Through extensive simulations and empirical applications on populations that have experienced multiple admixtures, we showed that gLike can accurately estimate dozens of demographic parameters when the true genealogy is known, including ancestral population sizes, admixture timing, and admixture proportions. Moreover, when using genealogical trees inferred from genetic data, we showed that gLike outperformed conventional demographic inference methods that leverage only the allele-frequency spectrum and yielded parameter estimates that align with established historical knowledge of the past demographic histories for populations like Latino Americans and Native Hawaiians. Furthermore, our framework can trace ancestral histories by analyzing a sample from the admixed population without proxies for its source populations, removing the need to sample ancestral populations that may no longer exist. Taken together, our proposed gLike framework harnesses underutilized genealogical information to offer exceptional sensitivity and accuracy in inferring complex demographies for humans and other species, particularly as estimation of genome-wide genealogies improves.

## Introduction

Accurately inferring the population history of humans has archaeological and historical significance, and it also helps to properly account for population structure in association studies and improve robustness in inferences about natural selection^1^. Because of the complicated interplay of random processes related to the underlying demography and observed genotypes – including migration, coalescence, recombination, mutation, and genotyping error – demographic inference is a challenging problem, often requiring simplifying assumptions or relatively coarse data summaries. One popular way of estimating the size changes of a single population utilizes a hidden Markov model (HMM) to describe the variation of haplotypes along the genome, where the hidden states correspond to the underlying genealogical trees^2–6^. As the number of potential trees grows exponentially with sample size, these methods are computationally scalable by tracking only a reduced representation of the underlying genealogy (e.g., SMC++^5^ and ASMC^6^ only track the coalescent times between a specific pair of haplotypes, while the remaining samples assume auxiliary functions). These methods are typically constrained by small sample sizes (usually <100) and the assumption of a single, homogeneous population, although they are flexible with respect to the population size trajectories over time. To accommodate for larger sample sizes that are more informative of the recent human history as well as more complex demographic events such as splits, migrations, and admixture, alternative approaches to demographic inference rely on a further reduced representation of the genealogy, the allele frequency spectrum (AFS)^7–11^. Although convenient to compute, the AFS may not contain enough information to recover the history precisely^12–14^.

HMM- and AFS-based methods are based on observed genotypes or haplotypes. However, since neutral variation is related to demographic history entirely via the genealogical processes, the (unknown) genealogy arguably has a more direct relationship with the underlying demography than the downstream genotypes^15–17^. Moreover, the complete genealogy of a collection of samples, as represented by an ancestral recombination graph (ARG)^18,19^, has richer information than the AFS since it includes additional data not reflected in the allele frequencies, such as the correlated coalescent histories between segments of a chromosome. Therefore, a genealogy-based demographic inference method has the potential to leverage the flexible topological structure of the ARG in distinguishing complex demographic histories, especially those with multiple admixtures.

Here, we introduce a genealogical likelihood framework named gLike to compute the likelihood of an observed genealogical tree under a parameterized demographic history. The intuition behind gLike is that a genealogy in itself does not imply the assortment history of any of its lineages (*i*.*e*. which set of discrete population memberships a particular lineage has traversed over time), meaning that all possible cases have to be considered. Notably, this idea bears similarity to the recently proposed “local ancestry path” problem by Pearson and Durbin^20^, but instead of inferring the population membership distribution of each individual node, gLike aims to compute the total likelihood of all combinations. By defining a “state” as the population memberships of all lineages existing at a specific time, possible movements between states throughout the history can be summarized into a directed acyclic *G*raph of *S*tates (GOS). We develop a full methodology for the GOS around three key problems: 1) constructing a minimal GOS that contains all necessary states; 2) computing the conditional probabilities between connected states with considerations of migrations, coalescences, and non-coalescences; and 3) propagating the marginal probabilities through the GOS to achieve the total likelihood of the tree, which can then be combined across multiple independent trees across the genome. As a general-purpose statistical framework and as a first step towards utilizing the information from the entire ARG, gLike is applicable to a variety of demographic events – migrations, splits, admixtures, and population size variations, providing tools for model selection and parameter estimation.

We demonstrate the advantage of genealogy-based demography inference by applying gLike to simulated scenarios, with particular emphasis on complicated admixture histories such as three- or four-way admixtures. gLike consistently outperforms existing AFS-based methods by producing parameter estimates closer to the simulated truth. In analyses of genotyped samples from Latino Americans and Native Hawaiians, the complex demography inferred by gLike is consistent with the known history of both admixed populations and their ancestral populations – Africans, Europeans, East Asians, Indigenous Americans, and Polynesians. Most notably, our inference required no reference sample from the ancestral populations (such as samples from Polynesians), nor explicit inference of local ancestries – information that is often not available or is imprecisely estimated for understudied populations with complex history.

## Results

### Method overview: genealogical likelihood under multi-population demography

A genealogical tree, despite being a complete record of the coalescent events of the sample haplotypes within a chromosomal interval, does not specify the migration history of lineages. In a typical genetic study, the samples (leaf nodes) are collected from known populations, which serves as the initial condition. The internal lineages could migrate, subject to the restriction that coalescences must happen within a population. Therefore, the probability of a given genealogical tree corresponds to the cumulative total of all migration scenarios that are compatible with this tree. Our proposed method, gLike, computes the likelihood of any given genealogical tree under a hypothesized demographic history (**Methods**). Operationally, it is broken into two topological steps to search for possible population memberships of lineages, followed by three numerical steps to compute the conditional and marginal probabilities (**Figure 1**).

**Fig 1.**
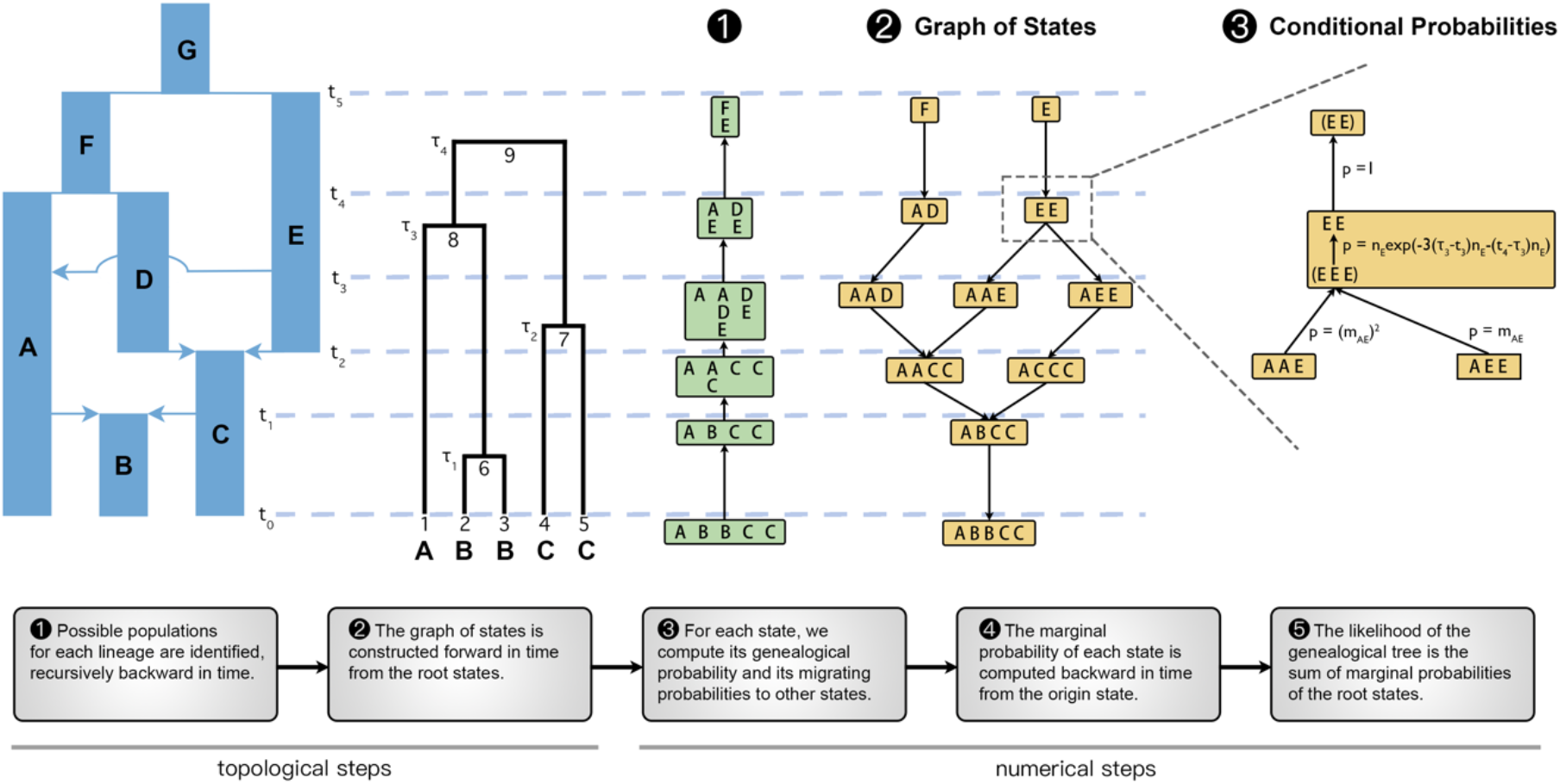
A schematic of the major steps of the gLike algorithm with examples. Starting from a parameterized demography and an observed genealogical tree with known sample populations, the fundamental data structure in gLike is the graph of states that summarizes all possible scenarios for all lineages to move through the populations across history. We denote the unique state at time zero that contains the observable population memberships of samples as the “origin state” (state “ABBCC” in this example), and the states about the root of the genealogical tree as the “root states” (states “F” and “E” in this example). The graph of states is constructed in Step 2, guaranteed by a preparatory Step 1 such that no redundant states will be generated, minimizing computational burden. Each column represents the population membership (in Step 2; e.g. “AD” means that lineage 8 is in population A and lineage 7 is in population D, *t*_4_ generations ago) or the set of possible memberships (in Step 1; e.g. at *t*_4_, lineage 8 may be in A or E, and lineage 7 may be in D or E,) of a certain lineage. In Step 3, the conditional probabilities are computed for all states in the GOS except the origin states, including the coalescence and non-coalescence probabilities implied in each state and the migration probabilities between connected states. Conditional probabilities are exemplified within the fourth epoch (between *t*_3_ and *t*_4_) around the state “EE”. Specifically, “EE” implies a unique hidden state “EEE” near the *t*_3_ end of the epoch because lineages 1 and 6 should both be in population E in order to coalesce into lineage 8, which is in E given the state “EE.” The connection between “EE” and “EEE” is represented by the “genealogical probability,” which consists of the probability that lineages 1, 6 and 7 did not coalesce before τ_3_ (with probability exp(−3(τ_3_ − *t*_3_)*n*_*E*_)), that lineages 1 and 6 coalesced at τ_3_ (with probability *n*_*E*_), and that lineages 7 and 8 did not coalesce before *t*_3_ (with probability exp(−(*t*_4_ − τ)*n*_*E*_)). The state “EE” has two child states, “AAE” and “AEE,” according to Step 2, connected via the intermediate state “EEE”. The transition from “AAE” to “EEE” requires two lineage migrations from “A” to “E,” which occurs with “migration probability” 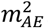. Similarly, transition from “AEE” to “EEE” occurs with probability *m*_*AE*_. In Step 4, the “marginal probability” of a state is defined as the probability conditional on the origin state and is computed recursively. For state “EE”, *p*(state *EE*) = *n*_*E*_ exp(−3(τ – *t*_3_)*n*_*E*_ − (*t*_4_ − τ)*n*_*E*_) ((*m*_*AE*_)^2^*p*(state *AAE*) + *m*_*AE*_*p*(state *AEE*)6. The marginal probabilities are propagated backward in time until the root states, and the log likelihood of the genealogical tree (conditional on the hypothesized demography) is, in step 5, the sum of all root states: *p*(tree) = *p*(state F) + *p*(state *E*).

We define a “state” as a specification of the population memberships of all lineages existing at a specific time. All possible states before each historical event (occurring at *t*_1,_ *t*_2_, …, *t*_5_ in this example) form a directed acyclic graph (step 2, **Figure 1**), which we call the “graph of states (GOS)”, a complete representation of all possible migration scenarios. When a state specifies a lineage in an impossible population, it becomes a dead-end state that does not connect to the origin. For example, in step 2, if we imagine a state “AA” at *t*_4_ as a child of “F”, it will not connect to the origin state “ABBCC”, because the fourth and fifth samples cannot migrate from C to A per the hypothesized demographic model (**Figure 1**; see also **Figure S1**). To reduce computation time, we avoid generating any dead-end state by a preliminary step (step 1, **Figure 1**) that summarizes possible population memberships for each lineage. For example, in step 1 at *t*_4_, lineage 8 may be in “A” or “E”, and lineage 7 may be in “D” or “E”, thus “AA” is not a legal state in step 2 (**Figure 1**). The graph of states is then constructed from the root states (“F” or “E” in this example) forward in time, by searching for child states according to both the specified migration events in the demography and the results in step 1. See **Figure S1** for intermediate results and further operational details during these two steps.

After building the GOS, the relevant conditional probabilities are computed. Because lineages are restricted to their respective population until a historical event, a state immediately before a historical event *t*_*s*_ is sufficient to specify the population memberships of all lineages between *t*_*s*−1_ and *t*_*s*_. For example, the state “EE” implies that not only the two lineages, but also the subtrees under both lineages are all in population E between *t*_3_ and *t*_4_. Given memberships of all lineages within the context of a state, we can compute the “genealogical probability” of the state based on standard coalescent theory to describe the coalescence (or lack thereof) events during the relevant interval on the tree. We also compute the “migration probability” between a state and its child, which is the product of the migration probability of each lineage, according to the migration matrix of the historical event (step 3, **Figure 1**). The “marginal probability” of a state is then the probability conditional on the origin state and can be computed recursively (step 4, **Figure 1**). Finally, we compute the likelihood of the genealogical tree as the sum of the marginal probabilities of the root states (step 5, **Figure 1**). See **Figure 1** legend for more explanation of genealogical, migration, marginal, and total probabilities related to the state “EE” in steps 3-5.

In practice, we apply gLike to a subsample of trees that are presumed independent, ideally from evolutionarily neutral sites distantly spaced across the genome (usually 10-100, depending on the computational resources), and the total likelihood is computed as the product over each individual tree. The total likelihood as a function of the demographic parameters is then optimized by simulated annealing. The final estimation of parameters is averaged over a number of subsamples with replacement. The variance across subsamples serves as an indicator of the uncertainty of the estimate.

### gLike accurately estimates all parameters in a three-way admixture demography

Admixed populations, especially those with three ancestral components or more, pose challenges to existing demographic inference methods. To showcase the performance of gLike to analyze complex admixture, we simulated 1000 haplotypes on a 30Mb chromosome from a population formed by two consecutive recent admixture events from three ancestral populations. Such a demography is parameterized by 3 event times, 2 admixture proportions, and 7 population sizes, totaling up to 11 parameters (**Figure 2A**). When true genealogical trees were available, the maximum likelihood estimates from gLike, averaged over 50 independent simulations, for all 11 parameters achieved an overall 3.8% relative error (**Figure 2B**), while gLike on the tsdate-reconstructed trees achieved an overall 23.3% relative error (**Figure 2C**).

**Fig 2.**
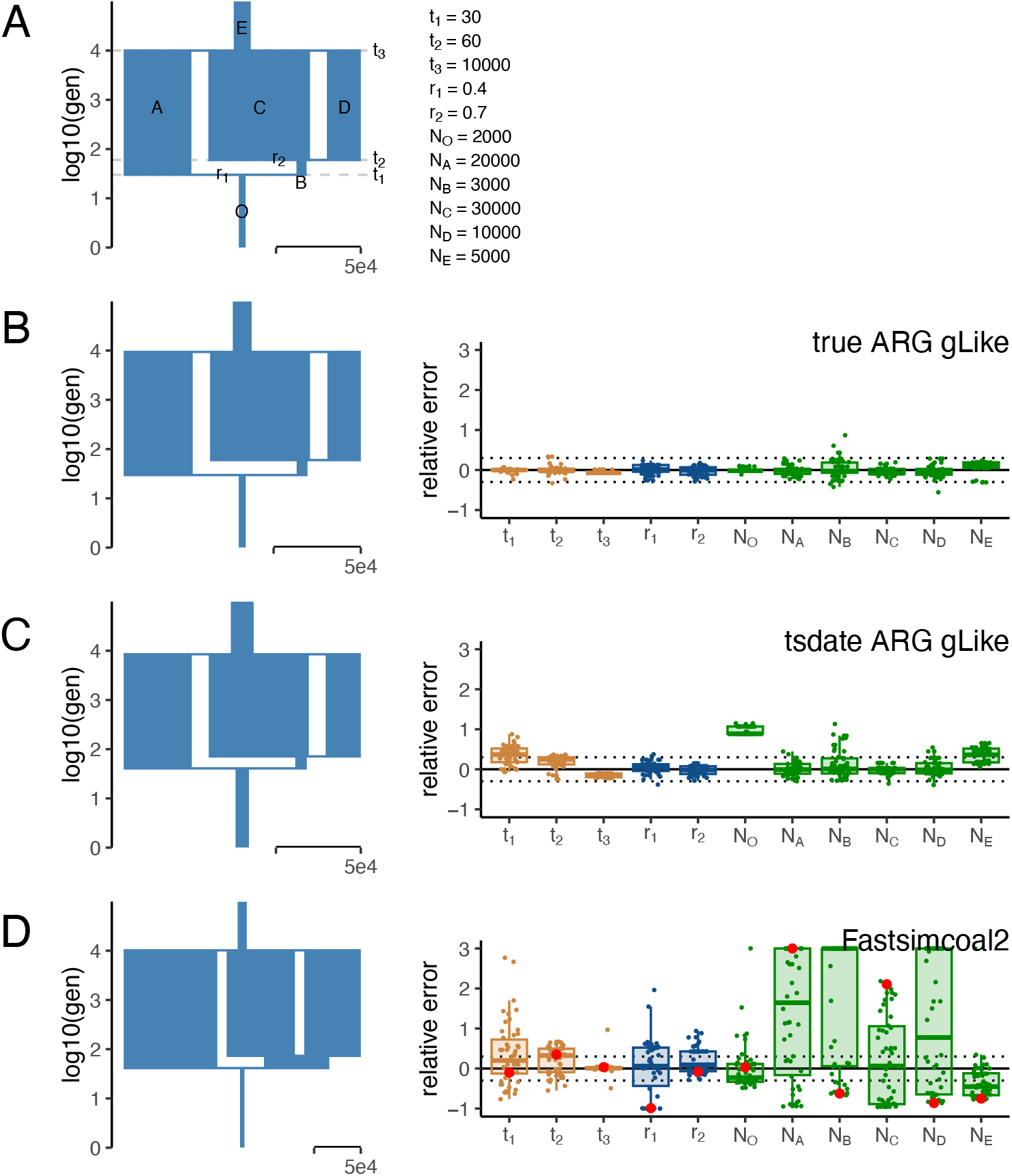
gLike accurately reconstructs three-way admixture without ancestral population samples. (A) The true demography under which the genealogical trees and genotypes were simulated, with 6 populations involved: population O is admixed from A and B; B is the intermediate population admixed from C and D, where C is defined to be the major ancestor (proportion ≥ 0.5) without loss of generalizability; E is the ancestor of A, C and D. All population sizes are to scale. There are 11 parameters involved, including 6 population sizes and: t_1_, time of admixture of population O; t_2_, time of admixture of population B; t_3_, time of split from population E; r_1_, admixture proportion of A in O; r_2_, admixture proportion of C in B. The true value of each parameter is provided on the right. (B-D) The reconstructed demography using parameter estimates averaged over 50 independent simulations (left) and boxplots of relative errors ((estimated-true)/true) in each simulation (right). Boxplots are capped at 300% relative error for ease of visualization. Trees and genotypes of 1000 haplotypes drawn from population O were simulated on a 30 Mb chromosome. The demographic parameters were estimated by gLike on the true trees (B), by gLike on the tsinfer+tsdate reconstructed trees from the true genotypes (C), and by Fastsimcoal2 on the allele frequency spectra derived from true genotypes (D). For Fastsimcoal2 results, the parameter estimates for the single run with the highest likelihood out of 50 independent runs, a practice commonly adopted by Fastsimcoal2, are labeled in red. A reference for the width of the population sizes equivalent to 50,000 is given in each panel.

We found that t_1_ and N_O_ are the most overestimated parameters (by 35.6% and 97.3%, respectively) when using tsdate-reconstructed trees, likely due to tsdate’s tendency to overestimate times of recent coalescences, prolonging the recent branches (**Figure S2**). Apart from t_1_ and N_O_, the other 9 parameters are estimated with 13.7% relative error.

We also tested Fastsimcoal2 (ref.^11^), which is capable of flexibly inferring complex demography using allele frequency spectra. Based on true genotypes and the same three-way admixture model, Fastsimcoal2 estimates had a relative error of 51.4%, which led to a visually distorted demography (**Figure 2D**). This is in sharp contrast to Fastsimcoal2 showing comparable accuracy to gLike on a three-population split demography (**Figure S3**). gLike also outperformed a Generative Adversarial Network (GAN)-based deep learning approach, pg-gan^21^, which was designed to overcome the limitations of relying on summary statistics such as the frequency spectrum. In our benchmarking, pg-gan performed well for a two-population split demography but was less accurate compared with Fastsimcoal2 and gLike on the three-population split and admixture demographies (**Figure S4** and data not shown). We thus did not test pg-gan further in this study. Nevertheless, our experiments with pg-gan were conducted without specialized neural network hardware and do not dismiss GANs’ potential as an emerging approach. Further training and improved procedures may enhance GAN-based demographic inference^22^.

We find that in our application with gLike for the demographies we have studied, analyses using tsinfer+tsdate-estimated genealogical trees produced more accurate estimated demographies those using trees estimated by Relate. The difference in performance may trace to the fact that recent coalescence times are overestimated by Relate to a greater extent than by tsdate, causing a 20∼50% depletion of coalescences within the recent dozens of generations (**Figure S2A**), thereby leading to mis-estimations in the gLike framework. As a result, gLike on Relate-reconstructed trees was not tested further in this study. Notably, Relate is more accurate in estimating the ancient part of the ARG, including the tree-wise times to the most recent common ancestor (tMRCAs) than tsinfer+tsdate (**Figure S2B**), which explains why in other applications utilizing the genealogical trees, such as inferring the genome-wide expected relationship matrix^17^ (eGRM), Relate may outperform tsdate.

### gLike detects components of admixture with high confidence

We examined the ability of gLike to distinguish two-way from three-way admixtures. We expect that the estimated parameters should reduce a complex model into a simpler one if the simpler model is closer to the true underlying model. Conversely, the likelihood should increase substantially when switching from a simple model to a complex one if the complex model is closer to the true underlying model. We first applied gLike under a hypothesized three-way admixture model to simulated trees and observed the estimated admixture proportions, r_1_ and r_2_ (**Figure 3A**, left and middle panels). Across 50 replicate simulations, when the true demography was a three-way admixture, the estimated admixture proportion for the third ancestry component, r_2_, centered around the true value (0.7) and was always far from the boundaries (0.5 and 1.0). When the true demography was a two-way admixture, the estimated r_2_ was almost always 1.0, with only one exception (**Figure 3A**). This indicates that gLike correctly reduced a three-way admixture model into a two-way model if the truth were indeed two-way admixed. In contrast, both r_1_ and r_2_ were estimated to be the boundary values around half of the time by Fastsimcoal2, regardless of the true demography (**Figure 3A**, right panel).

**Fig 3.**
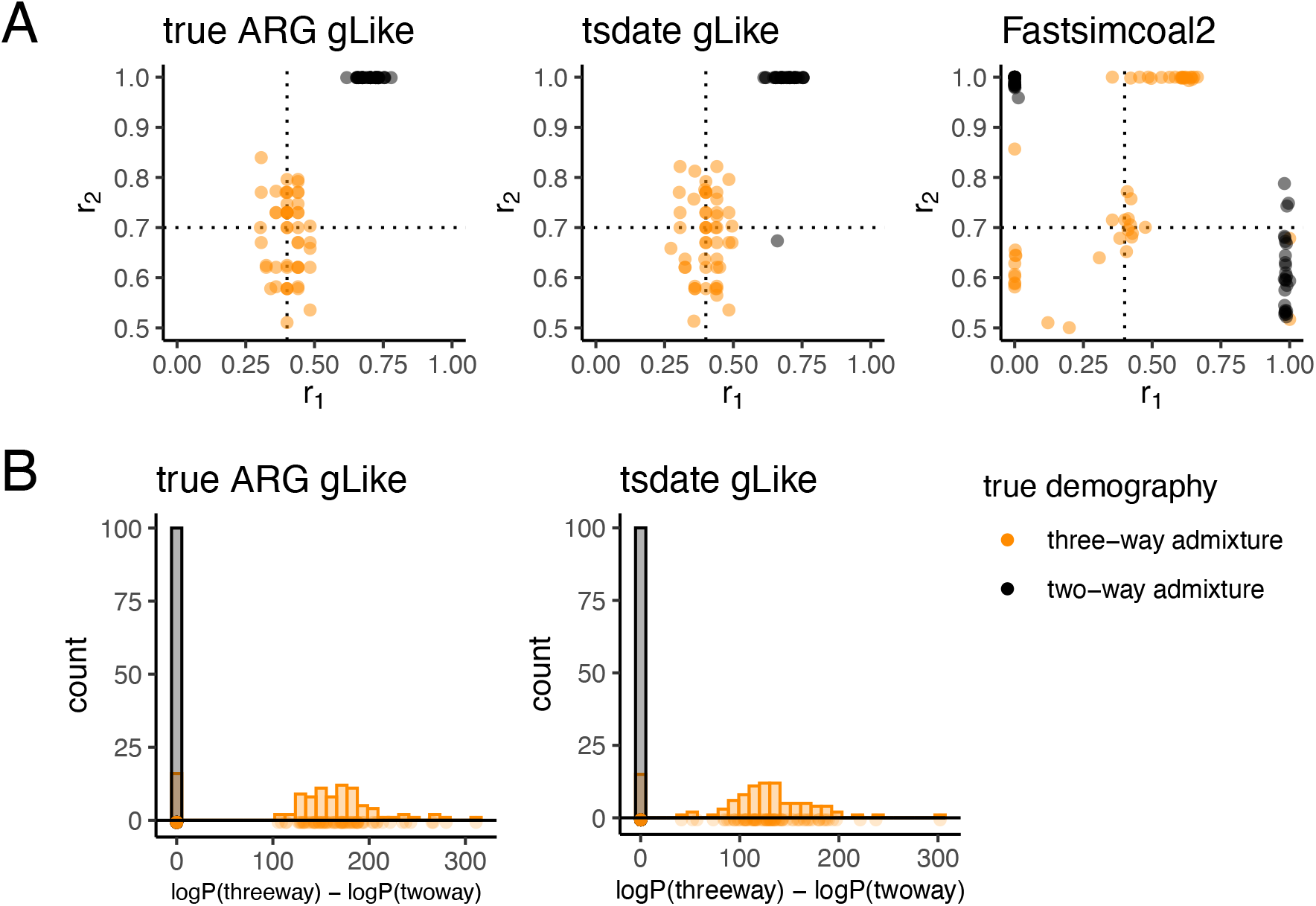
gLike distinguishes three-way admixture from two-way admixture. True (left) and tsifner+tsdate reconstructed (middle) trees were obtained from simulated three-way (orange, same model as **Figure 2**) and two-way (grey, r_2_ was set to 1, removing contribution from population D) admixed populations. (A) gLike was applied assuming a three-way admixture model. The estimated r_1_ and r_2_ values in each of 50 independent simulations are shown, dashed lines denote true values of r_1_ and r_2_ in three-way admixture simulations. (B) gLike was first applied under a two-way admixture model, then the model is expanded into a three-way admixture and gLike likelihood is optimized while fixing shared parameters between two models (see **Methods** for technical details). The distributions of log likelihood improvement after model expansion are shown as histogram. Model selection through the Akaike information criterion (AIC) resulted in a classification accuracy of 92%.

We next evaluated the maximum likelihood achieved under a two-way admixture model and a three-way admixture model (**Methods**). AIC model selection was applied on the log-likelihood differences between two models to select the more likely model between the two-way and three-way admixtures. Across 100 independent simulations, the three-way admixture model was never preferred when the true admixture was two-way, and the three-way admixture model was preferred over two-way when it was the true model ∼85% of the time with both true ARGs and tsdate-reconstructed ARGs, resulting in a ∼92% accuracy of classification.

### gLike reproduces complex demographic histories from stdpopsim

Having established that gLike sensitively detects components of admixtures and estimates parameters with high accuracy, we further evaluate its ability to reconstruct two additional demographic models with increasing complexity, as published in stdpopsim^23^ – the American Admixture (stdpopsim model 4B11; **Figure 4**) and the Ancient Europe (stdpopsim model 2A21; **Figure 5**) demographies.

**Fig 4.**
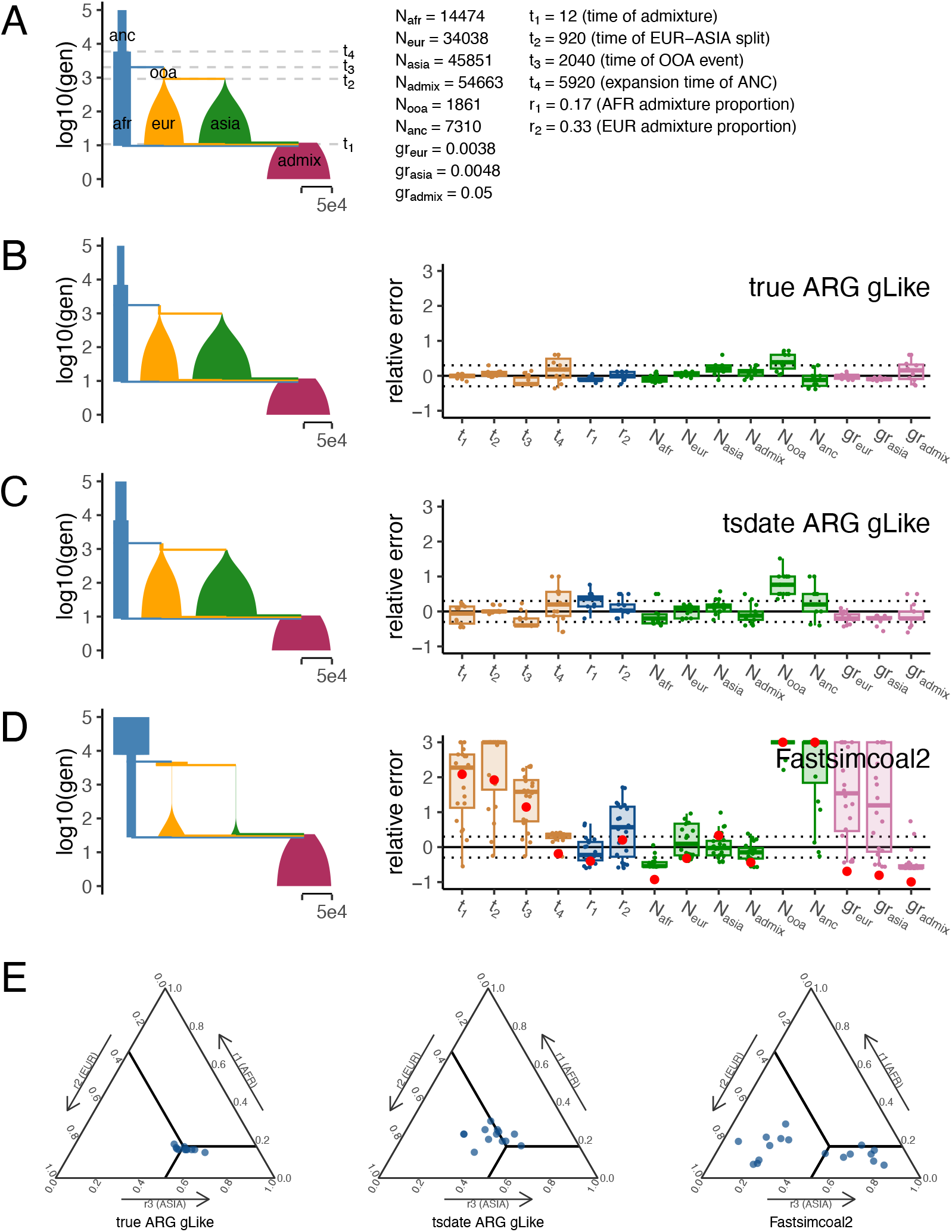
gLike reconstructs the American admixture demography. (A) American admixture demography with parameters from stdpopsim model 4B11. All population sizes are drawn to scale. (B-D) The reconstructed demography using estimations averaged over 50 replicate simulations (left) and boxplots of relative errors in each simulation (right). Trees and genotypes of 1,000 haplotype from the admixed population were simulated on a 30 Mb chromosome, the demographic parameters were estimated by gLike on the true trees (B) or the tsinfer+tsdate reconstructed trees (C), and by Fastsimcoal2 on the allele frequency spectra derived from true genotypes (D). Boxplots are capped at 300% relative error for ease of visualization. For Fastsimcoal2 results, the parameter estimates for the single run with the highest likelihood out of 50 independent runs are labeled in red. A reference for the width of the population sizes equivalent to 50,000 is given in each panel. (E) Ternary plots showing admixture proportions estimated by gLike on the true trees (left), by gLike on the tsinfer+tsdate reconstructed trees (middle) or by Fastsimcoal2 on the allele frequency spectra of the true genotypes (right), with slide lines indicating true parameters.

**Fig 5.**
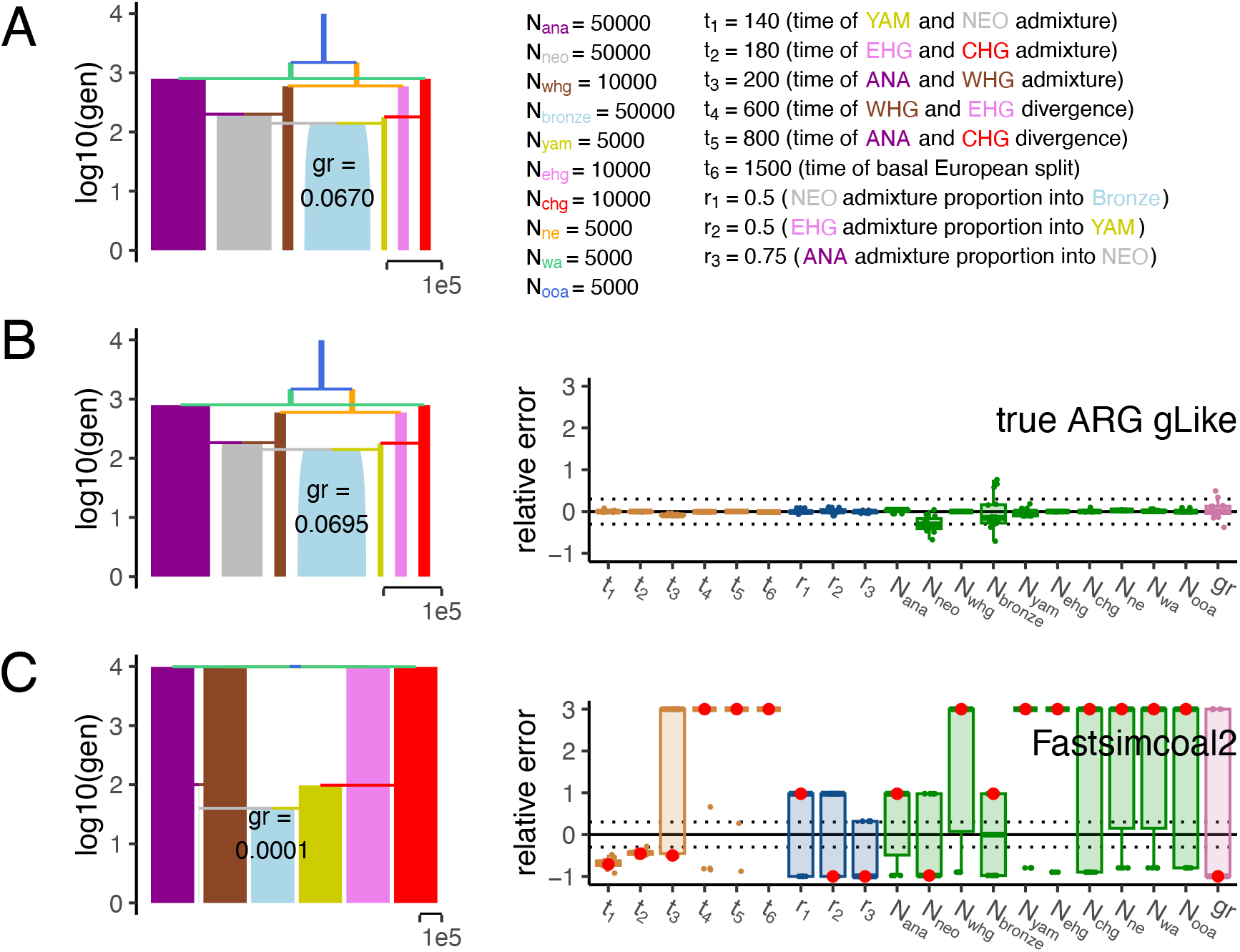
gLike reconstructs the ancient Europe demography. (A) Ancient Europe demography with parameters from stdpopsim model 4A21. The Bronze Age population is plotted with initial size true to scale, but the growth rate is shown as text to avoid a disproportionate figure. All other population sizes are constant size and drawn to scale. (B, C) The reconstructed demography using estimates averaged over 50 replicate simulations (left) and boxplots of percentage errors in each simulation (right). Trees and genotypes were simulated on a 30 Mb chromosome. A total of 2200 haplotype samples (1000 contemporary samples descended directly from the Bronze Age population and 200 ancient samples each from the six ancient populations) were drawn at collection times as described by stdpopsim. The demographic parameters were estimated by gLike on the true trees (B) or by Fastsimcoal2 on the allele frequency spectra of the true genotypes (C). Boxplots are capped at 300% relative error for ease of visualization. For Fastsimcoal2 results, the parameter estimates for the single run with the highest likelihood out of 50 independent runs are labeled in red. A reference for the width of the population sizes equivalent to 10,000 is given in each panel.

The American Admixture model consists of four populations (AFR, EUR, ASIA and ADMIX; **Figure 4A**), where ADMIX is formed by a very recent admixture from the other three populations. This model has 15 parameters, including 4 event times, 2 admixture proportions, 6 population sizes and 3 exponential growth rates. We simulate 1000 haplotypes from population ADMIX on a 30Mb chromosome. gLike on the true trees inferred all 15 parameters with overall 11.3% relative error (**Figure 4B**). The majority of the error was in N_ooa_, the size of the out-of-Africa predecessor of the European population, which was overestimated by 38.5%. gLike on the tsdate-reconstructed trees inferred parameters with overall 23.5% relative error (**Figure 4C**). Except from the overestimation of N_ooa_ by 77.8%, the error concentrated on the African branch. For example, r_1_ (the African admixture proportion) was overestimated by 30.2%, and N_anc_ was overestimated by 27.1%. Fastsimcoal2, in comparison, estimated the same set of parameters with 258.7% relative error (**Figure 4D**). Fastsimcoal2 estimated the African proportion fairly accurately, but appears unable to distinguish between the European and Asian proportions (**Figure 4E**).

As AFS-based methods presumably have better performance in the presence of a multi-dimensional allele frequency spectrum, we compared gLike and Fastsimcoal2 in additional simulations where 500 haplotypes from each ancestral population were sampled to supplement the 1000 admixed samples (**Figure S5)**. Presence of ancestry reference samples improved the accuracy and consistency of Fastsimcoal2’s estimation of almost all parameters (an average of 213.1% relative error), especially the admixture proportions. But gLike based on the true and inferred trees (5.8% and 16.7% relative errors, respectively) was still more accurate in capturing the histories of these populations (**Figure S5**).

To test gLike’s performance on intra-continental admixtures, we also evaluated the Ancient Europe model from stdpopsim (2A21). This model is a four-way admixture model where the two intermediate ancestors of Bronze Age population are each in turn admixed from two ancestors (**Figure 5A**). We simulated 1000 haplotypes from the present-day population that descended from the Bronze Age, and 200 from each of the ancient populations, according to the times specified by stdpopsim. Applying gLike to the true trees resulted in estimates of the 20 parameters with overall 3.0% relative error (**Figure 5B**). The main misestimated parameter was the 29.6% underestimation of N_neo_, an ancient population that only existed for 20 generations (180-200gen) when its samples were collected. Fastsimcoal2 estimated all parameters with an average relative error of 132.3% (**Figure 5C**). The estimates of several population sizes reside near the preset borders -- a behavior that has been suggested to be an intrinsic pitfall of AFS-based methods^24^. We did not test tsdate in this experiment because its ARG inference method does not currently make full use of the ancient samples (instead, they are inserted as “proxy sample ancestors” onto the existing ARG). Given our evaluation above, however, we would expect that gLike will substantially improve over Fastsimcoal2 in accuracy of parameter estimates if inferred ARGs can accurately incorporate ancient samples, and that gLike can generally handle intra-continental admixtures when ancestral populations may be relatively closely related.

### Inferring admixture history of Latinos and Native Hawaiians using genome-wide array data

We applied gLike to investigate populations with complex demographic history using genome-wide genotyping data from Latinos and Native Hawaiians, each with 500 subsampled diploid individuals. We parameterized a four-way admixture model consisting of Africans, Europeans, East Asians and a fourth ancestral population, which is used to model the Indigenous Americans (for Latinos) or the Polynesians (for Native Hawaiians). We estimated genealogical trees from the genotyping data using tsdate and estimated a total of 16 parameters using gLike (**Figure 6**; **Supplemental Table 1**). We estimated the Latino lineages to be 10.7% from Africans, 44.2% from Europeans, 45.1% from Indigenous Americans, and 0% (across all 20 independent threads) from East Asians, while the Native Hawaiian lineages were 19.8% from Europeans, 33.4% from East Asians, 46.8% from Polynesians, and 0% (across all 20 independent threads) from Africans (**Figure 6**). As expected, we estimated the Native Hawaiians to be more recently admixed than the Latinos (19 compared to 25 generations ago). Also, the Native Hawaiians had a slightly smaller initial population size than the Latinos (35,682±10,656 compared to 41,579±16,851; but both are likely overestimated. See **Discussion**) and grew at a slower rate (0.078±0.009 compared to 0.132±0.012) since the admixture.

**Fig 6.**
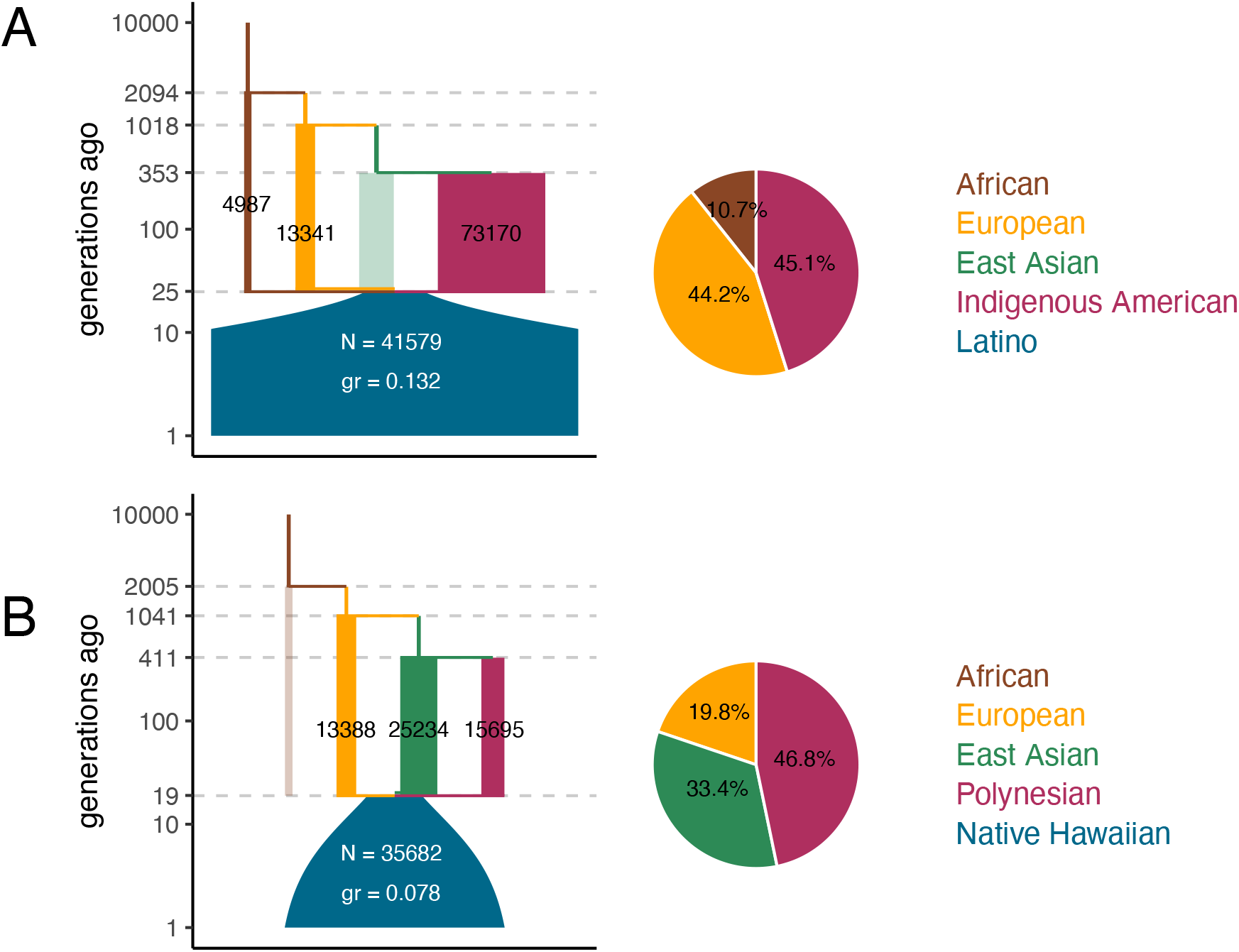
parameter estimations for the demographic histories of Latinos and Native Hawaiians. gLike was applied under a potential four-way admixture model reminiscent of stdpopsim model 4B11 for both the Latino (A) and Native Hawaiian (B) data. The four potential ancestral populations are African, European, East Asian, and Indigenous American (for Latinos) and Polynesian (for Native Hawaiians). The reconstructed demographic diagrams are to scale, marked with relevant parameters. N, size of the admixed population in diploids at time of admixture; gr, growth rate of the admixed population. Ancestral populations estimated to have 0% admixture proportion are shown as translucent, because their sizes cannot be estimated. Pie charts show the estimated admixture proportions of ancestral populations.

The European ancestries participated in both admixtures. As expected, we found the estimates of its population size (13,388±2,388 and 13,341±4,702) and of time of divergence with the East Asians (1,018±172 and 1,041±87 generations ago) to be highly concordant between two data sets, suggesting the same underlying population that colonized the Americas and Polynesia. Note that this ancestry should be more appropriately interpreted as the ancestral population responsible for the colonization, which is less genetically diverse than the entire European continent currently or at the time. The Indigenous Americans and Polynesians, though represented as the same component in the model, were estimated to have different sizes (73,170±28,939 compared to 15,695±7,393), which may reflect greater population sizes or more extensive structure in the ancestors to the Latino samples than to the Native Hawaiian samples. Considering the potential errors during the ARG-reconstruction process (as have been seen in **Figures 2, 4** and **S2**) and biases due to the lack of high-quality sequencing data for these two admixed cohorts (**Table S2**), these estimates of the demographic parameters for both populations should be taken with caution. Nevertheless, our results suggest that gLike is able to qualitatively capture known features of the demographic history of Latinos and Native Hawaiians without reference data from their ancestral populations, and the results stand to improve as ARG-reconstruction approaches advance.

## Discussion

With the fast development of scalable ARG inference over the past few years, development of population-genetic approaches that explicitly use the ARG or its marginal trees is an exciting area of active research. With this in mind, our current study introduced a framework that explains the stochastic formation of the genealogical trees in a multi-population context, and computes the full likelihood of each demographic scenario. Our results revealed that the history of at least three ancestral populations can be clearly decoded from the genealogical trees of a single admixed sample without knowledge of the ancestral populations. For many understudied diverse populations across the world, it is often unclear whether they are admixed, and if so, what the ancestral populations were. Even if the ancestral populations are known or can be hypothesized, they likely no longer exist or are difficult to sample. For these populations, demographic inference using allele frequencies is difficult, since distinct demographic scenarios can give similar AFSs^12^. gLike has the potential to provide new insights into studies of these understudied or ancient populations, as well as the demographic history of other species.

It is worth clarifying that the admixture proportions in the demographic context (such as those estimated by Fastsimcoal2^11^ and gLike here) have a slightly different meaning from that in the genomic context (such as those estimated by STRUCTURE^25^ and ADMIXTURE^26^). As a demographic parameter, the admixture proportion describes the probability of a lineage to migrate (backward in time) from one population to another, while in the genomic context, this proportion describes how much of the genome one population shares with another. The two concepts can deviate primarily in two cases: 1. There is considerable genetic drift after the admixture, especially when the population size is small; 2. The admixed population, O, may have a genetic component from, say, population A, not because A participated in the formation of O, but because of more ancient migrations from A to other ancestries of O. In gLike results, all admixture proportions should be interpreted in demographic context. In practice, admixture proportions could be estimated through other means in the genomic context, and then be used as the initial values for gLike to improve optimization speed and stability, while allowing gLike to make further adjustments as needed.

We also note that currently gLike is not utilizing the full information encoded in an ARG, but rather is relying on sets of presumed independent trees. In many ways, gLike was inspired by HMM-based demographic inference^2–6^, where genealogical trees are implicitly utilized. However, these methods are computationally intensive and have limited scalability, primarily due to the intricate handling of recombination events. We reasoned that while recombination events are essential for ARG inference, they are less informative for ARG-based demographic inference. Once the ARG (and thus the genealogical trees within the ARG) has been accurately inferred from the genotypes, reliance on recombination events for insights into demography becomes less important. Recombination can be modeled as a random breakpoint in the genealogical tree re-coalesced onto the rest of the tree – the random break is independent of demography, and the re-coalescence holds minimal information compared to the numerous coalescences already on the tree. In light of this, gLike currently focuses on rigorously modeling lineage assortments and coalescent events within individually independent trees, rather than the variability between neighboring trees, to achieve greater scalability (in order to handle thousands of samples and multiple populations). Future enhancements of gLike may then model recombination to incorporate the remaining information encoded in the ARG. Furthermore, gLike has some commonality with approaches to species-tree inference based on gene trees, where gene trees can be used to estimate the topology and branch lengths of a phylogenetic tree^27^. Whereas such methods estimate the whole topology, we pre-specify the demographic history and estimate parameters related to it, including processes like admixture that do not feature as prominently in species-tree inference. In cases where the demographic history is sketchy, it may be possible to develop approaches akin to the species-tree inference to estimate parts of the topology.

One current limitation of gLike is that certain parameters are not individually identifiable, but could only be optimized in combination. For example, the effects of population size and growth rate are hard to separate if a population exists for only a short time (**Figure S6**). Any combination of the two parameters that produces the same average coalescence rate will have a similar likelihood, making it difficult to identify the global optimum. Such entangled parameters are in fact a limitation in many demographic inference methods and often result in similar likelihoods for many combinations of parameters. When applying gLike with hill-climbing-based optimization methods, the estimates of entangled parameters could be path dependent. Thus, a grid search on specific entangled parameters after a general optimization routine may be beneficial to an unbiased estimation of the demography.

In addition, continuous migration is not currently supported by gLike, because it drastically increases the number of states. In the American Admixture simulations (**Figure 4**), we omitted the weak migrations (10^−5^-10^−4^ per generation) between continental populations as originally specified by the stdpopsim model. Omitting the continuous migrations have no visible impact on estimating the remaining parameters unless they are ∼100 times more intense than that currently specified in the stdpopsim model and presumed to be typical between continental human populations (**Figure S7**). However, such frequent migrations (10^−3^-10^−2^ per generation) may exist between intra-continental populations where geographical separations are minimal. Estimating the migration rate itself is also of interest in ecological studies of other species, and a future focus will be extending gLike to incorporate continuous migration. One obvious solution is to discretize the continuous migration into a number of pulse migrations, which results in many layers each containing a large number of states. An effective discretization strategy, as well as an efficient random sampling technique on the states, seems necessary to address this challenge.

Current ARG inference methods have achieved remarkable scalability and accuracy, but their biases and errors still deserve attention in genetic applications. We have showcased the varying performance of tsinfer+tsdate and Relate at different time scales in admixed populations (**Figure S2**). The overestimation of branch lengths at recent times appears to be a common problem for both methods, but is more severe in Relate-inferred trees, to the degree that meaningful GOSs are difficult to construct. Tsinfer and tsdate are also faster because they use heuristic algorithms to avoid the O(n^2^) pair-wise comparisons. However, the bottom-up approach of tsdate is somewhat less accurate for ancient coalescences, whereas Relate’s hierarchical clustering-based method infers the deep part of the genealogies with higher accuracy (especially beyond 1000 generations ago), and thus captures global relatedness more robustly^17^. There may be techniques to adjust one’s result with the other, thus combining both of their advantages. With scalable and accurate ARG inference across broader scales, we expect the reliability and accuracy of gLike demographic inference to be further improved.

Finally, we acknowledge that human migrations and admixtures exist on a continuum. In the current framework we opted to model discrete populations and components of ancestries, as is customary when modeling the histories of recently admixed populations such as the Latinos.

But one of the advantages of an ARG-based view of human history may be to remove the notion of distinct populations. Enabling continuous rather than pulse-like migrations between populations to enhance gLike may be another step forward, but future developments of ARG-based demographic inference may emphasize on the paradigm shift to represent human histories and structure on a continuum.

## Methods

### Formalization of the problem: Probability of a genealogical tree under a demography

The demographic history of *K* populations can be represented by the interplay between two stochastic processes affecting the lineages – coalescence and movement among populations. The coalescence rate *n*_*a*_(*t*) of each population *a* as a function of time *t* is

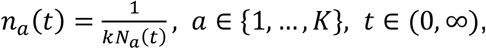

where *N*_*a*_ is the effective population size, and *k* is ploidy. And the migration probability matrix *m* at each of the *S* historical events is

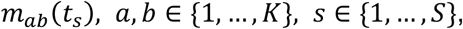

where *t*_*s*_ is the time of the *s*-th historical event, and *m*_*ab*_(*t*_*s*_) is the instantaneous probability for a lineage to move (backward in time) from population *a* to *b*.

The demography is thus defined as

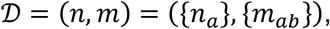

a size-*K* vector of coalescence rates defined on continuous time, and a *K* × *K* matrix of migration probabilities defined on a discrete set of times. While gLike currently does not explicitly incorporate continuous migration, it can potentially be represented as a series of historical events through discretization.

A genealogical tree with *N* nodes can be defined by the time and children of each node

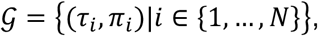

where τ_*i*_ is the time of the node *i* (or equivalently, the emergence of lineage *i*), and π_*i*_ is the set of its child nodes (which is empty if *i* is a leaf node). The end time ω_*i*_ of lineage *i* can be calculated as time of its parent node (that is, ω_*i*_ = τ_*j*_ if *i* ∈ π_*j*_) or ∞ if it has no parent. Our goal is to compute ℙ(𝒢|𝒟) for arbitrary 𝒢 and 𝒟, and we will omit thereafter the “conditional on 𝒟” notation, which is always implied.

It is helpful to define the set of lineages existing at time *t* as

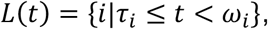

and the lineages emerging between *t* and *t*′ as

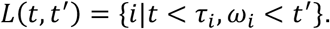

### Migration trajectory and states

The population identity of a lineage *i* during its existence,

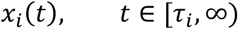

is a time-dependent variable taking values from {1, …, *K*}that describes how this lineage, or its ancestor lineage when *t* > ω_*i*_, migrates in history. For convenience, the value of *x*_*i*_ (*t*) at exactly the time of a historical event is defined as the left limit 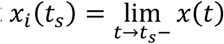, so that *x*(*t*) is left-continuous.

The population identity of all lineages existing at any time throughout the history is

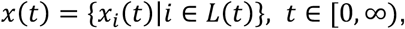

which gives a complete migration trajectory of the genealogical tree. The genealogical tree itself does not dictate *x*, and the probability of it should be computed as the sum over all possible trajectories,

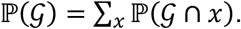

In order to compute *ℙ*(*𝒢*) recursively over time, we define *𝒢* (0, *t*) as the genealogical history in *𝒢* until time *t*, and define a “state” as

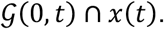

For example, the state “ABCC” in **Figure 1** at *t*_1_contains *𝒢* (0, *t*_1_), which indicates that lineages 2 and 3 coalesced at τ_1_but all other possible coalesces has not happened at *t*_1_, and *x*(*t*_1_) = ABCC, which indicates that the remaining four lineages (1,6,4 and 5) are in populations A,B,C and C, respectively, at *t*_1_.

Now *ℙ*(*𝒢*) can be expressed as the sum of probability of root states

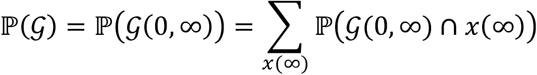

### Conditional probability between states

The conditional probability between states

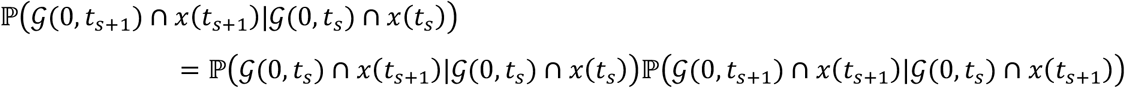

consists of a migration probability and a genealogical probability.

The migration probability

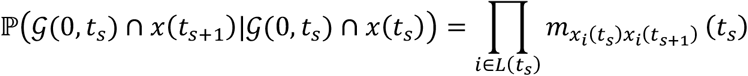

describes the migration of each lineage *i* from *x*_*i*_ (*t*_*s*_) to *x*_*i*_ (*t*_*s*+1_) at time *t*_*s*_.

The genealogical probability *ℙ*(*𝒢* (0, *t* _*s*+1_) ∩ *x*(*t* _*s*+1_)| *𝒢* (0, *t* _*s*_) ∩ *x*(*t* _*s*+1_)P describes how likely the genealogical tree grows according to *𝒢* backward in time from *t*_*s*_ to *t* _*s*+1_, given population identities *x*(*t*_*s*+1_). This requires that every coalescence in *G* happened exactly at its time in *G* (which we call the coalescence probability) and that any other possible coalescence did not happen (which we call the non-coalescence probability).

The coalescence probability is

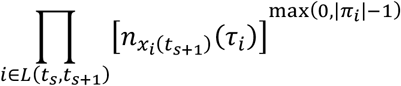

where 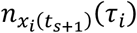 is the coalescence rate of lineage *i*’s population when it emerges. Note that the lack of migration between τ_*i*_ and *t* _*s*+1_ guarantees *x*_*i*_ (τ _*i*_) = *x*_*i*_ (*t* _*s*+1_). And max(0, |π_*i*_| − 1) is the number of coalescences at the emergence of *i* (for example, a binary node is formed with one coalescence, a ternary node can be viewed as two coalescences at the same moment, and a leaf node or unary node does not have coalescence).

The non-coalescence probability is

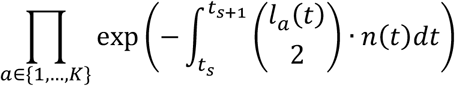

where

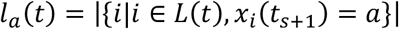

is the number of lineages in population *a* at time *t* (if population identities are specified by *x*_*i*_(*t*_*s*+1_)), which is a step function that jumps when lineages emerge or coalesce; 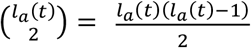 is the number of lineage pairs in *a* that are possible to coalesce; and the exponential term is the probability that none of them actually coalesced during (*t*_*s*_, *t*_*s*+1_), which is derived from a nonhomogeneous Poisson process with rate 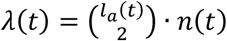. Note that *n*(*t*) can be any integrable function, enabling flexibility to the population size variation in the demographic model.

We conclude that the conditional probability between states is

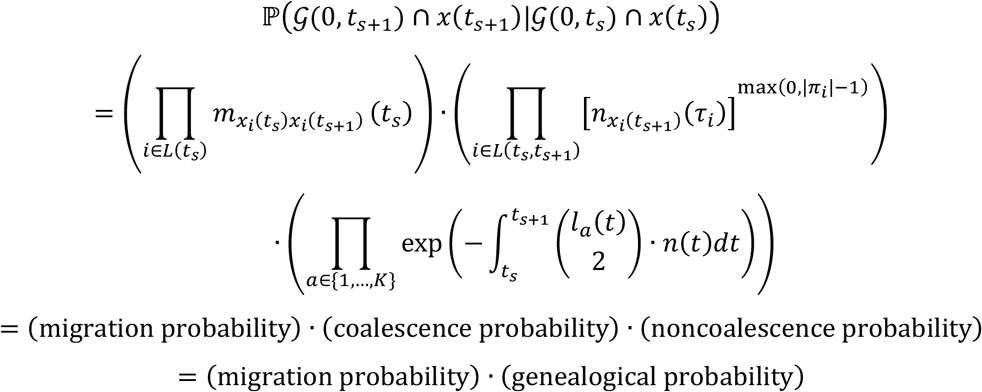

Practically, the migration probability has to be computed between any parent-child state pair, but the genealogical probability is independent from the child state and needs to be calculated only once for every state. As a boundary condition, the origin state at the bottom (*i*.*e*. leaves) of the tree has probability one

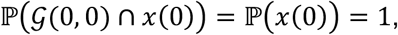

where *x*(0) specifies the population identities of each individual in the study samples.

### The minimal graph of states

All possible states at all times of all historical events *t*_1_, *t*_2_, …, *t*_*S*_ form a directed acyclic graph, named as the graph of states (GOS), where states in adjacent layers (one at *t* _*s*_ and the other at *t*_*s*+1_) are connected with their conditional probability as introduced above. A state with zero marginal probability will not contribute to the marginal probability of its parent state and is redundant in the graph. A GOS without redundant states is called a minimal GOS.

The coalescence probability and non-coalescence probability are always above zero, because population sizes cannot be zero or infinity. This means that, to judge if a state is possible or not, we only have to check the migration probabilities, which are decomposable into migrations of each individual lineage. In other words, a state is possible if every lineage is in a possible population. To put it mathematically, we have

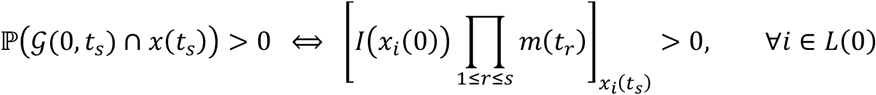

where *I*(*x*_*i*_ (0)) is a size-Kindicator vector with value 1at the population *x*_*i*_ (0) where sample *i* was collected, and all other elements zero; *∏*_1≤*r*≤*sm*_(*t*_*r*_) is the transition matrix summarizing the first *s* historical events; and 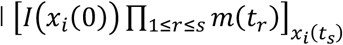 is the probability that lineage *i* migrated from *x*_*i*_(0) to *x*_*i*_(*t*_*s*_). **Figure 1** step 1 can be understood as the non-zero elements in *I*(*x*_*i*_ (0))*∏* _1≤*r*≤*s*_*m*(*t*_*r*_) for every *s*.

### Implementation details and optimization

With the above-mentioned theory to calculate *ℙ*(*𝒢*|*D*_*θ*_) on a demographic model *D*_*θ*_ parameterized by *θ*, the estimated parameters that best explains the observed *𝒢* is

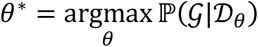

gLike encapsulates the likelihood computation and a simulated annealing-based optimization into an open-source Python package, alongside a C extension to accelerate Cartesian product operations when searching for child states (GitHub page: https://github.com/Ephraim-usc/glike). All probabilities are implemented in log scale, and sums of probabilities are calculated with the scipy logsumexp function. If the number of states at a layer exceeds the preset limit (10^*#*^ by default), a random subsample of states is generated to approximate the likelihood. When multiple, presumed independent and neutrally evolving, trees are provided, the final log likelihood is the sum of log likelihoods of each tree. We presume independence of trees as the total likelihood would assume more complicated forms if trees were nearby and not independent. We also presume neutrality as coalescence probabilities would deviate from the inverse of population sizes when there are variants under natural selection. We set a user-defined parameter to drop some proportion (default: 50%) the lowest likelihood trees during optimization, as we found in practice that this filtering improves robustness against errors in tree reconstruction (such as erroneous coalescences) and migrations that are neglected in the demographic model.

### Demographic inference in simulations

All simulations were performed on a 30 Mb chromosome with both recombination and mutation rates set to 10^−8^ per generation per base pair, with a sample size of 1,000 haplotypes from the admixed population. The demographic parameters are annotated in the corresponding figures, or cloned from stdpopsim^23^ models 4B11 (American Admixture) and 4A21 (Ancient Europe). In American Admixture simulations, we ignored the continuous migrations in our simulations and estimations. The extent to which hidden migrations potentially undermines gLike results was tested on additional simulations with 1-, 10- and 100-times continuous migrations as reported by stdpopsim 4B11. In the Ancient Europe simulation, we additionally sampled 200 haplotypes from each ancestral population according to the collection times reported by stdpopsim, in order to mimic genetic studies with ancient DNA.

To evaluate gLike, ARGs and genotypes were simulated by msprime^28^. ARG reconstructions by tsinfer+tsdate^29,30^ or Relate^31^ were performed with all default parameters as suggested in the user manual. One hundred evenly spaced trees across the chromosome were selected for gLike inference. The precision of gLike parameter estimation (i.e., the minimal step size during optimization by simulated annealing, relative to the current estimate) was set to 2%. The absolute difference between the average estimate and the truth, divided by truth, is defined as the relative error. The average estimates across 50 replicate simulations were used as the final pictorial representation of the reconstructed demography, with boxplots of the relative errors across 50 replicates also shown. The standard deviation across 50 replicate simulations serves as an indicator of the parameter uncertainties as listed in **Tables S1** and **S2**.

To compare gLike to Fastsimcoal2 (ref ^11^), derived allele frequency spectra were computed on all simulated SNPs (including singletons), and parameter estimation was performed with 100,000 simulations and 40 ECM (expectation/conditional-maximization) loops, using the commands “-n 1 -s0 -d -k 1000000” for AFS simulation and “-n 100000 -s0 -d -M -L 40” for parameter estimation. The estimate with the highest likelihood obtained among 50 independent runs was used as the final pictorial representation of the reconstructed demography (following the same practice recommended by the authors of Fastsimcoal2^32^), with estimates from all 50 shown in the accompanying boxplots. We also compared gLike performance to pg-gan^21^, a deep learning demographic parameter inference method that uses generative adversarial networks to create realistic simulated training data. Genotypes from simulated ARGs of the same demographic model were used as training data, run for up to 300 training iterations with default training parameters. We also used the same range for each demographic parameter to be consistent with the Fastsimcoal2 comparisons. Since pg-gan gives multiple sets of parameter proposals at end of training, the set of inferred demographic parameters with the lowest relative error compared to the true parameters was selected as the final estimate of this run. A total of 50 independent runs were conducted.

To characterize the impact of ARG reconstruction using array data instead of sequencing data, we performed additional simulation experiment in which SNPs were retained with the probability

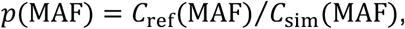

where MAF is the minor allele frequency of the simulated SNP, *C*_*r*ef_(*MAF*) is the number of occurrences of MAF in the Latinos array data, and *C*_*s*im_ is the number of occurrences of MAF in a simulated genome (3,000Mb). As expected, it was found that *C*_*s*im_ is greater than *C*_*r*ef_ across all values of MAF ∈ [0, 0.5], which ensures p is always less than one. We then inferred the ARG using tsinfer+tsdate using the simulated array data.

### Model selection in simulations

To test for the existence of an additional ancestral component, gLike was applied under a two-way admixture model and a three-way admixture model, and the maximum likelihoods achieved under both models were compared. Specifically, the two-way admixture model structurally mimicked the three-way admixture as in **Figure 2A**, but without population D, so that all lineages from population B entered population C. As such, the two-way admixture model had two fewer parameters – r_2_ (admixture proportion from D) and N_D_ (population size of D). gLike was then applied in a two-step manner. First, the parameters were estimated under the two-way admixture model with the default hill-climbing optimization. Next, we applied gLike under the three-way admixture model and perform a grid search on r_2_, N_C_ and N_D_, while fixing other parameters at their two-way admixture estimates. Finally, the difference between the maximum log likelihoods achieved under two models was used for AIC model selection (with 2 degrees of freedom, to account for the two extra parameters in the three-way admixture model), and the model with the higher AIC value was selected.

### Latinos and Native Hawaiians data processing

A total of 5,382 self-identified Native Hawaiians and 3,659 self-identified Latinos from the Multiethnic Cohort (MEC) were genotyped on two separate GWAS arrays: Illumina MEGA and Illumina Global Diversity Array (GDA). After taking the intersection of SNPs found on both arrays, the genotyping data were lifted to hg38 using *triple-liftover*^33^ to ensure alleles in inverted sequences between reference genome builds were properly lifted. We removed variants that were genotyped in fewer than 95% of individuals, variants out of Hardy-Weinberg Equilibrium (p < 10^−6^), and individuals with greater than 2% missing genotypes (though no one was removed with this threshold). After quality check, the Native Hawaiian and Latino datasets contained 990,549 and 1,093,693 SNPs, respectively. The data were phased without a reference using EAGLE^34^ and its default hg38 genetic map. We randomly subsampled 1,000 haploids and removed monomorphic SNPs, resulting in 879,040 and 927,254 SNPs in the Native Hawaiian and Latinos datasets, respectively. The ancestral alleles were called by a comparison with the human ancestor GRCh38 e107 genome (URL: ftp.ensembl.org/pub/release-86/fasta/ancestral_alleles/). Tsinfer and tsdate were used with all default parameters as suggested in the user manual to reconstruct the ARG. The human neutralome^35^ (*i*.*e*., the regions of the human genome identified as likely selectively neutral) was converted into hg38 coordinates, and 319 neutral regions that are at least 5Mb from each other were selected for gLike analysis. Ten trees were sampled in each gLike optimization thread, and 20 threads were run in parallel. The estimates of demographic parameters were averaged over 20 threads. The precision of gLike parameter estimation was set to 5%, higher than 2% used in simulations. This choice is due to the broader span of the likelihood curve’s plateau, which generally extends beyond 5%, wider than observed in simulations. Therefore, using smaller step sizes would increase computational costs with little gain in performance.

## Data Availability

The individual level genetic data for Native Hawaiian and Latino datasets were derived from the Multiethnic Cohort (MEC), and are available on dbGaP (accession numbers: phs000220.v2.p2 and phs002183.v1.p1). The gLike package is available on its github page (https://github.com/Ephraim-usc/glike2).

## Acknowledgement

We would like to thank Iain Mathieson and Sara Mathieson for discussions and advice. Research reported in this publication was supported by National Institute of Health under award number R35GM142783 and R01HG12605 to C.W.K.C. Computation for this work was supported by USC’s Center for Advanced Research Computing (https://carc.usc.edu).

## Author’s Contributions

C.W.K.C., D.O.D.V., and C.D.H. conceived of the study. C.F. and C.W.K.C. designed the study. C.F. and J.L.C. performed the analysis. B.L.D. curated the data. C.F., M.D.E., N.A.M., and C.W.K.C. interpreted the data. C.F., J.L.C., M.D.E. and C.W.K.C. wrote the manuscript with input from all co-authors.

