## Supplemental Figures for "A likelihood-based framework for demographic inference from genealogical trees"

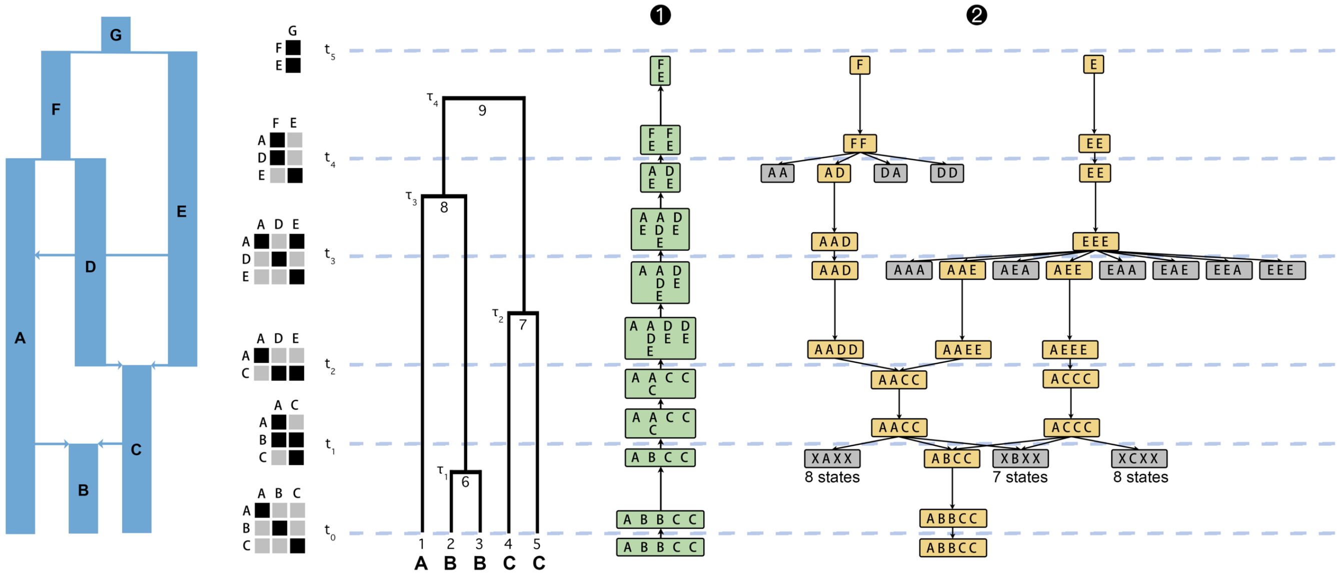


**Figure S1. A detailed look at the topological steps.** Intermediate results in the first two steps of Figure 1 are shown to illustrate the technical details. The non-zero elements in the mass migration matrices in each of the historical events are marked as black, and zeros are marked as grey. Results in Step 1 and Step 2 are shown at the beginning and end of each phase. Each column represents the population membership (in Step 2) or the set of possible memberships (in Step 1) of a certain lineage. Step 1 proceeds backward in time from the bottom. Note that the possible populations of the parent lineage are the intersection of the possible populations of its children (for example at $\tau_3$, lineage 1 can be in populations A or E, while lineage 6 can be in populations A, D or E. Because coalescence could only happen in the same population, the parent lineage 8 can be in populations A or E, but not D). Step 2 proceeds forward in time from the root states – "E" and "F" in this case. Crossing a historical event, the candidate children states of a state is the Cartesian product of the possible destination populations of each lineage. The candidate children that conflict with the results in Step 1 are dead-end states marked in grey (they are shown for clarification purpose only and are not generated in practice). For example, in step 2, If we imagine a state "AA" at $t_4$ as a child of "F", it will not connect to the origin state "ABBCC", because the fourth and fifth samples cannot migrate from C to A per the hypothesized demographic model. Note that the historical event at $t_5$ is older than the root of the genealogical tree at $\tau_4$ and has no effect during the whole process.

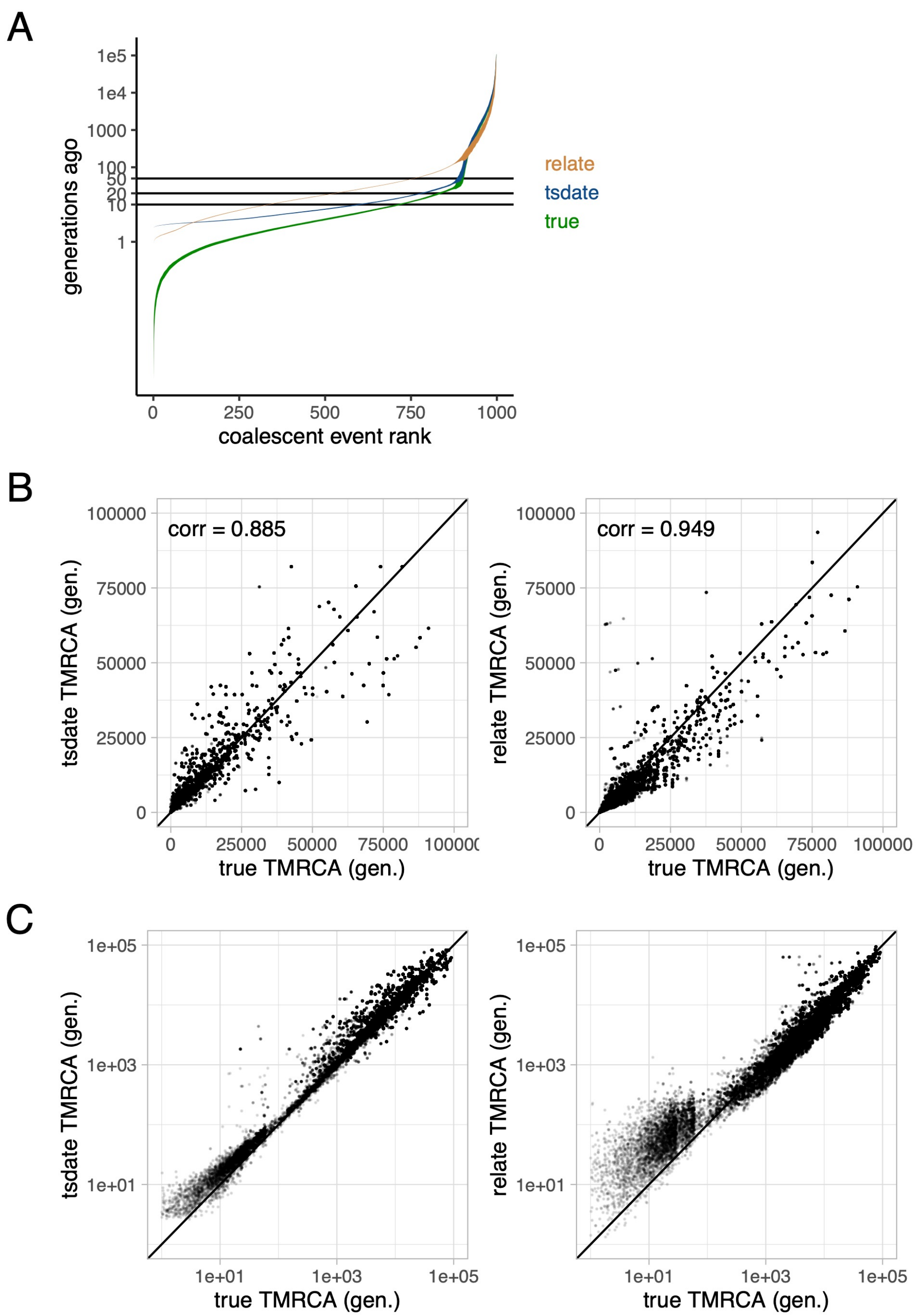


**Figure S2. Bias of tsinfer+tsdate and Relate on admixed data.** (A) The times of coalescences (i.e., inner nodes) in ascending order in a three-way admixed genealogical tree with 1000 haplotypes, ribbons showing 2 times standard deviation across 50 independent simulations. The true trees were simulated under the three-way admixture demography as in **Figure 2A**.
(B) TMRCA in the true tree versus tsinfer+tsdate (left) and Relate (right) reconstructed tree. Results from 50 independent simulations are shown together. (C) Same data as in (B) but shown in log scale. Note the substantial bias of relate TMRCA under 100 generations.

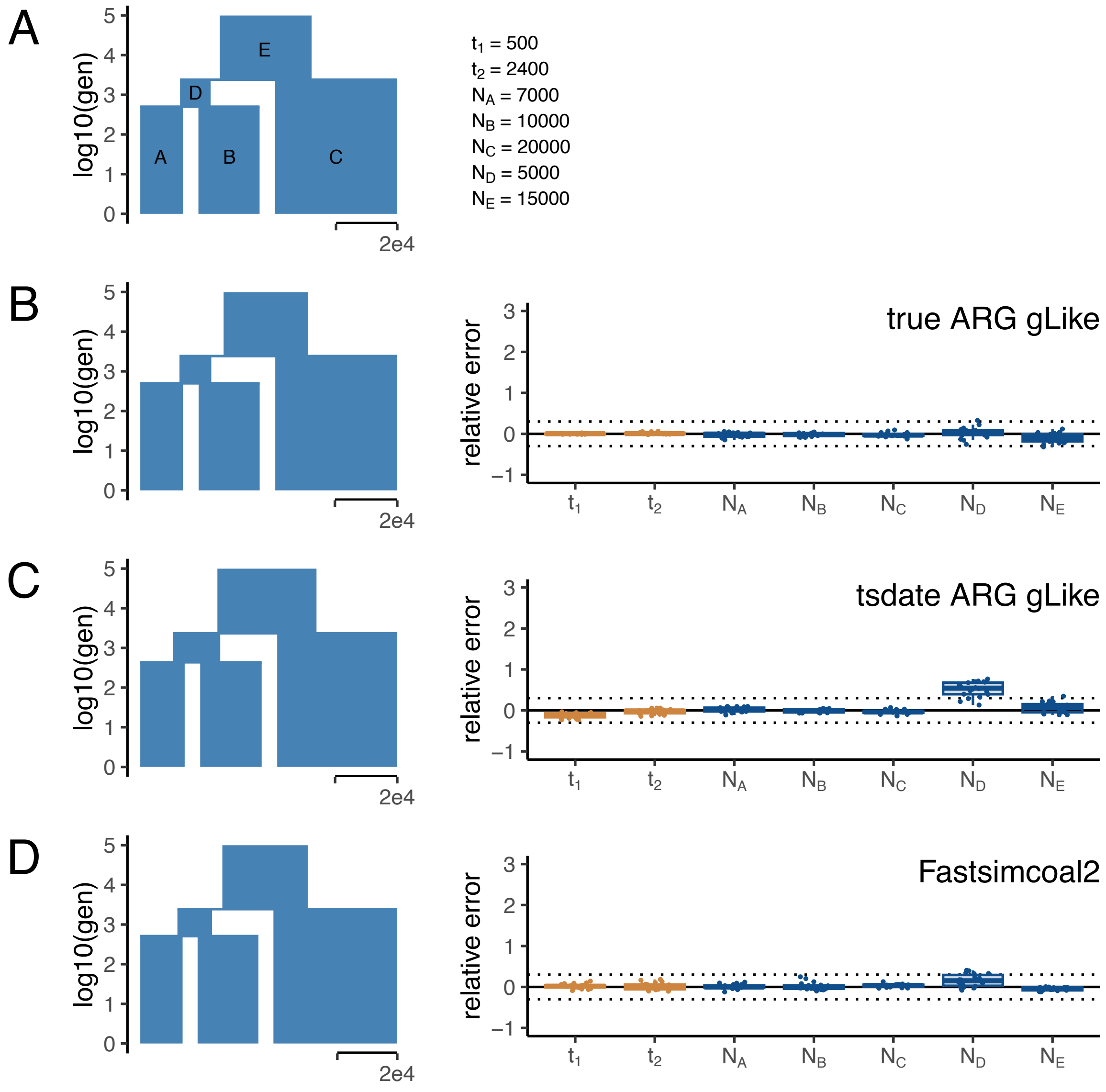


**Figure S3. gLike and Fastsimcoals perform comparably on three-way split demography.** (A) The true demography in which the ancestral population E splits into C and D at $t_2$, and D splits again into A and B at $t_1$. The true value of each parameter is provided on the right. (B-D) The reconstructed demography using parameter estimates averaged over 50 independent simulations (left) and boxplots of percentage errors in each simulation (right). ARGs and genotypes were simulated on a 30 Mb chromosome, with 500 haplotypes drawn from each of populations A, B and C. The demographic parameters were estimated by gLike on the true ARGs (B), by gLike on the tsinfer+tsdate reconstructed ARGs (C), and by Fastsimcoal2 on the allele frequency spectrum derived from true genotypes (D). Without admixture events, Fastsimcoal2 and gLike performed comparably in terms of accuracy for parameter estimations.

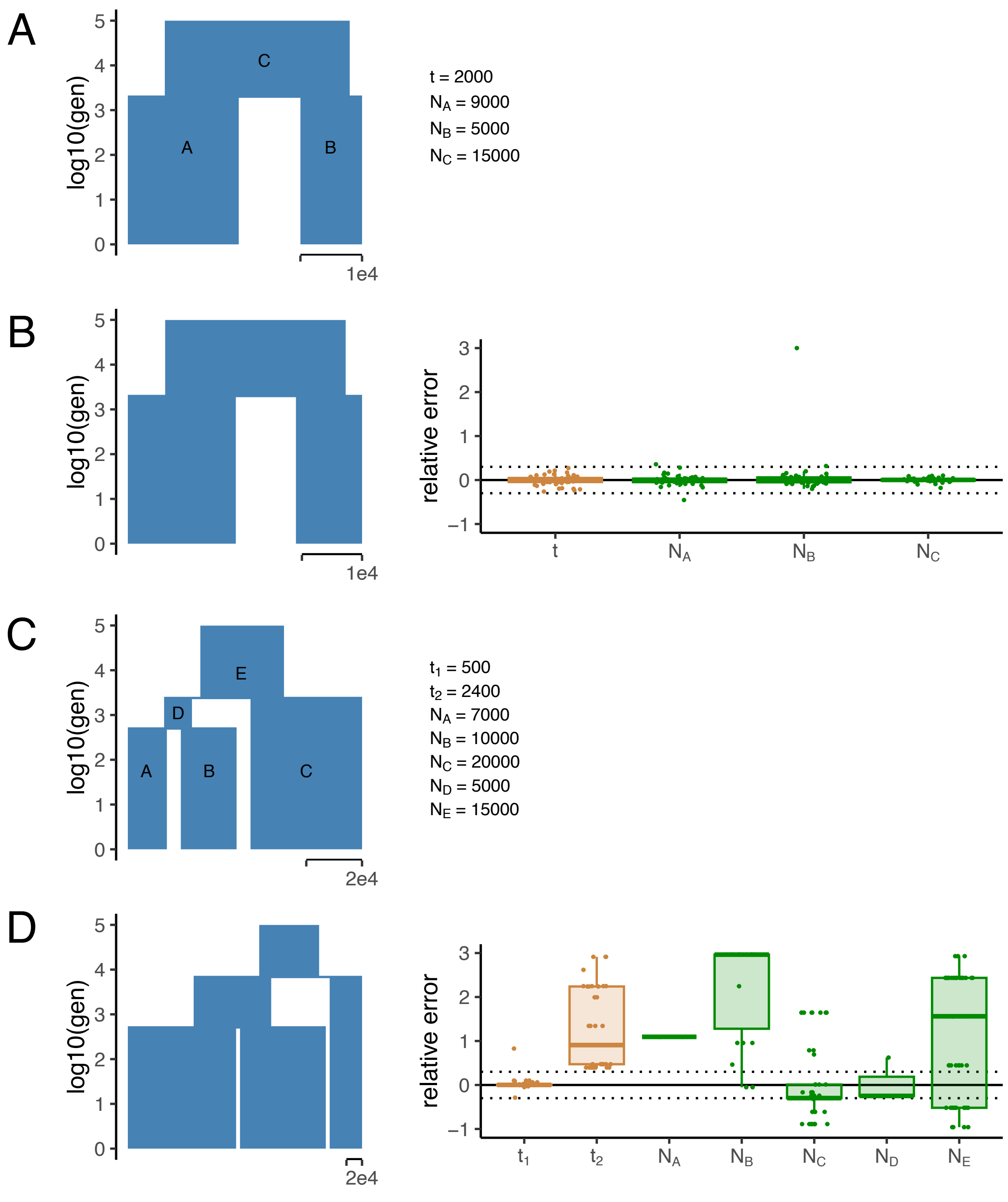


**Figure S4. pg-gan fails to infer three-way split demography.** Genotypes are simulated under the two-way spilt (A) and three-way split (C) demographies. The true value of each parameter is provided on the right. The demographic parameters were estimated by pggan on the true genotypes. (B and D) The reconstructed demography using parameter estimates averaged over 50 independent simulations (left), and boxplots of percentage errors in each simulation (right). ARGs and genotypes were simulated on a 30 Mb chromosome, with 500 haplotypes drawn from each population. While pg-gan accurately reconstructed the two-way split demography, its performance on the three-way split demography was notably worse.

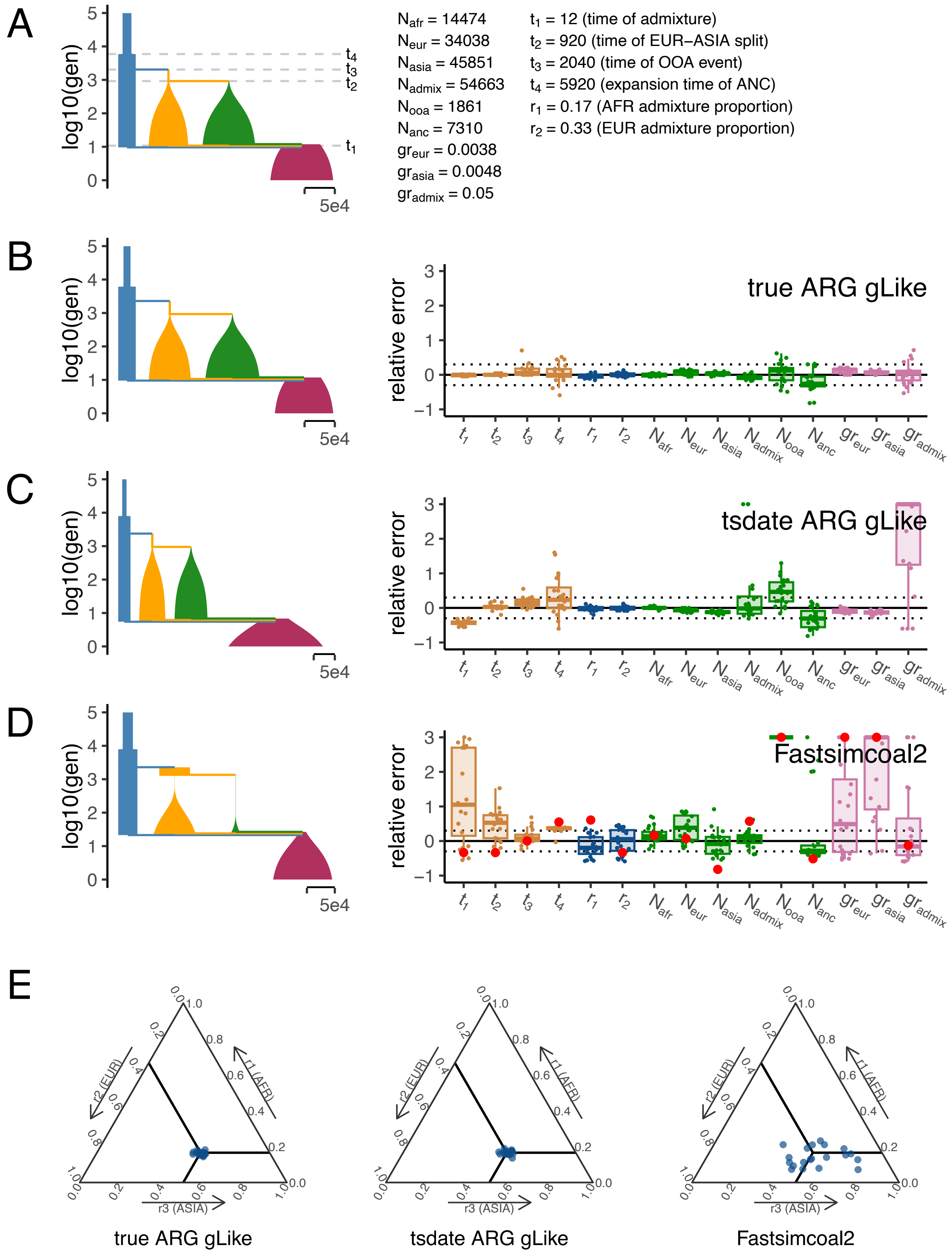


**Figure S5. Reconstructing the American admixture demography with reference samples.** The same experiment in **Figure 4** except that 500 haplotypes from AFR, EUR and ASIA are also collected, totaling up to a sample size of 2,500 including the 1,000 admixed haplotypes. Panels (A-E) are organized similarly as **Figure 4**. For Fastsimcoal2 results, the parameter estimates for the single run with the highest likelihood out of 50 independent runs are labeled in red. Having ancestral reference samples improved the estimates for admixture proportions for gLike with tsinfer+tsdate reconstructed tree, but did not generally noticeably improved the performance with fastsimcoal2.

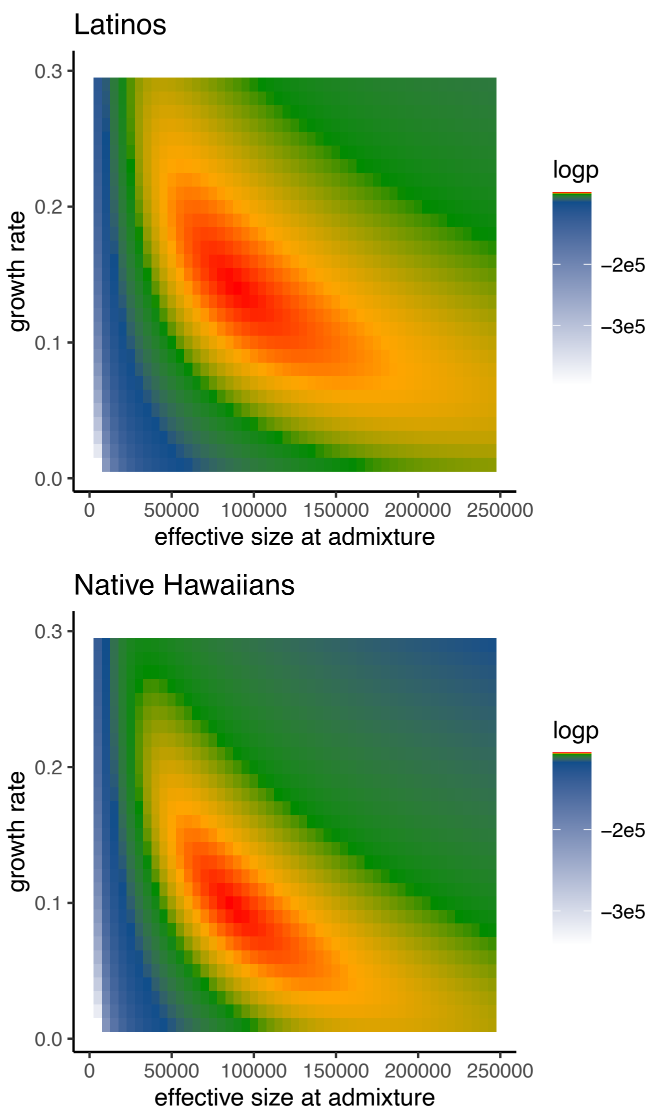


**Figure S6 . Unidentifiability between population sizes and growth rates.** The log-likelihood of the gLike model on the population sizes (at time of admixture) and growth rates of the Latinos and Native Hawaiians in a grid of possible parameters. All other parameters were fixed at their estimates shown in **Figure 6**. This result indicates the potential bias when estimating entangled parameters, because the hill-climbing optimization could stop anywhere along the red curve, depending on the initial values.

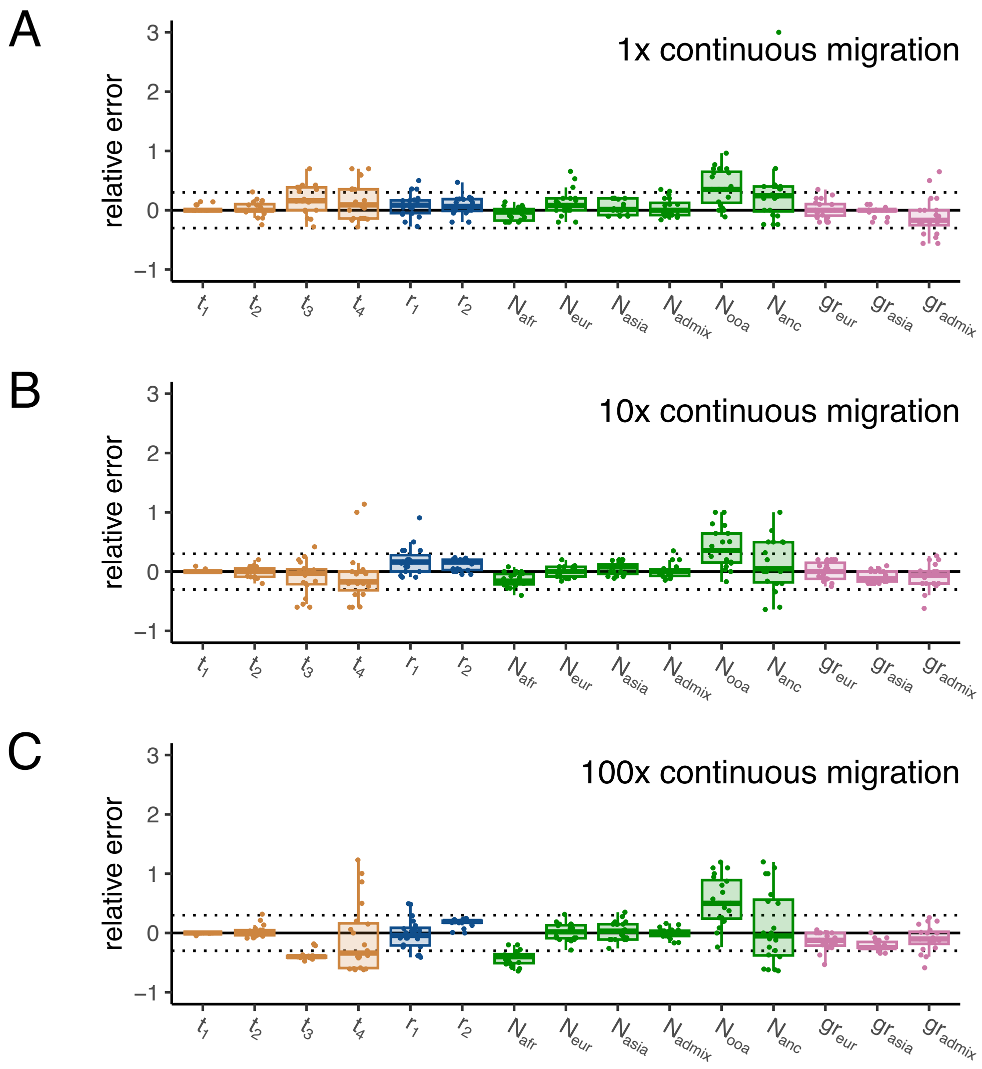


**Figure S7. Robustness of gLike against misspecified continuous migrations.** The same experiment in **Figure 4A** except that the true demography contains AFR-EUR, AFR-ASIA, EUR-ASIA and AFR-OOA continuous migrations that are set to be 1x (A), 10x (B) and 100x (C) of their rates as in the stdpopsim 4B11 model. gLike was applied on the true trees in the same way as in **Figure 4A**, assuming no continuous migrations. Note that the 1x continuous migrations have no visible impact on the results, while 100x continuous migrations lead to considerable underestimations of $t_3$, $t_4$ and $N_{\mathrm{afr}}$, due to the accumulation of coalescences earlier than expected in a migration-free demography.
