## Supplemental Tables for "A likelihood-based framework for demographic inference from genealogical trees"

| | Description | Latinos mean ± std | Native Hawaiians mean ± std |
|---|---|---|---|
| $t_1$ | Time of admixture (gen) | 24.8 ± 9.1 | 18.9 ± 2.3 |
| $t_2$ | Time when indigenous Americans or Polynesians split from Asians (gen) | 353 ± 49 | 411 ± 42 |
| $t_3$ | Time when Asians split from Europeans (gen) | 1018 ± 172 | 1040 ± 87 |
| $t_4$ | Time when Europeans split from Africans (gen) | 2094 ± 326 | 2004 ± 58 |
| $r_1$ | African admixture proportion | 0.107 ± 0.068 | 0.0 |
| $r_2$ | European admixture proportion | 0.442 ± 0.148 | 0.198 ± 0.012 |
| $r_3$ | Asian admixture proportion | 0.0 | 0.334 ± 0.036 |
| N | population size at admixture | 41579 ± 16850 | 35682 ± 10656 |
| $N_{afr}$ | African population size | 4986 ± 426 | NA |
| $N_{eur}$ | European population size | 13341 ± 4701 | 13388 ± 2388 |
| $N_{asia}$ | Asian population size | NA | 25234 ± 6984 |
| $N_{ia/pol}$ | indigenous Americans or Polynesians population size | 73170 ± 28939 | 15695 ± 7392 |
| $N_{aa}$ | Asian population size between $t_2$ and $t_3$ | 3092 ± 958 | 2702 ± 795 |
| $N_{ooa}$ | European population size between $t_3$ and $t_4$ | 2948 ± 612 | 2470 ± 558 |
| $N_{anc}$ | African population size before $t_4$ | 2846 ± 716 | 2665 ± 444 |
| gr | Growth rate of admixed population (per gen) | 0.132 ± 0.012 | 0.078 ± 0.009 |

**Table S1. Latinos and Native Hawaiians parameters estimates and uncertainties.** Estimated quantities corresponding to the results in **Figure 6**. Uncertainty was calculated as the standard deviation cross 20 independent threads, each thread containing 10 distant and selectively neutral trees (see **Methods**). Population size estimates marked as NA are not estimable, because the admixture proportion from such population was estimated to be zero.

| | Sequencing data gLike mean ± std | Array data gLike mean ± std | Relative difference |
|---|---|---|---|
| $t_1$ | 40.7 ± 6.9 | 32.4 ± 2.4 | -20.3% |
| $t_2$ | 71.3 ± 9.3 | 55.5 ± 12.3 | -22.2% |
| $t_3$ | 8474 ± 483 | 7321 ± 391 | -13.6% |
| $r_1$ | 0.418 ± 0.056 | 0.425 ± 0.039 | 1.6% |
| $r_2$ | 0.680 ± 0.079 | 0.693 ± 0.065 | 1.9% |
| N | 3946 ± 194 | 3212 ± 48 | -18.6% |
| $N_A$ | 20078 ± 3364 | 16434 ± 1820 | -18.1% |
| $N_B$ | 3398 ± 1006 | 2932 ± 1417 | -13.7% |
| $N_C$ | 29484 ± 3163 | 21871 ± 3069 | -25.8% |
| $N_D$ | 10328 ± 1866 | 7879 ± 1376 | -23.7% |
| $N_E$ | 13534 ± 4654 | 10892 ± 2404 | -19.5% |

**Table S2. Bias in inferred parameters from gLike using tsdate-inferred trees from simulated array vs. sequencing data.** The same three-way admixture demography as **Figure 2A** was simulated. Genotypes were subsampled from simulated sequencing data to match the empirical MAF distribution in the Latinos genotyping data (**Methods**). tsdate was used to infer the ARG based on either the simulated genotype data or the sequencing data. Genealogical trees from each ARG were sampled and analyzed by gLike to compare the bias in parameter estimates due to using only a subset of variations typically found on an array.
